# Development of a NanoBRET assay for evaluation of 14-3-3σ molecular glues

**DOI:** 10.1101/2023.12.31.573792

**Authors:** Holly R. Vickery, Johanna M. Virta, Markella Konstantinidou, Michelle R. Arkin

**Author notes:** Corresponding Author Michelle R. Arkin – Department of Pharmaceutical Chemistry and Small Molecule Discovery Center, University of California, San Francisco 94143, United States, Address: 600 16^th^ St. GH S512D San Francisco, CA 94143, United States. H.R.V. and J.M.V. contributed equally to this work.

## Abstract

We report the development of a 384-well formatted NanoBRET assay to characterize molecular glues of 14-3-3/client interactions in living cells. The seven isoforms of 14-3-3 are dimeric hub proteins with diverse roles including transcription factor regulation and signal transduction. 14-3-3 interacts with hundreds of client proteins to regulate their function and is therefore an ideal therapeutic target when client selectivity can be achieved. We have developed the NanoBRET system for three 14-3-3σ client proteins CRAF, TAZ, and estrogen receptor α (ERα), which represent three specific binding modes. We have measured stabilization of 14-3-3σ/client complexes by molecular glues with EC_50_ values between 100 nM and 1 μM in cells, which align with the EC_50_ values calculated by fluorescence anisotropy in vitro. Developing this NanoBRET system for the hub protein 14-3-3σ allows for a streamlined approach, bypassing multiple optimization steps in the assay development process for other 14-3-3σ clients. The NanoBRET system allows for an assessment of PPI stabilization in a more physiologically relevant, cell-based environment using full-length proteins. The method is applicable to diverse protein-protein interactions (PPIs) and offers a robust platform to explore libraries of compounds for both PPI stabilizers and inhibitors.

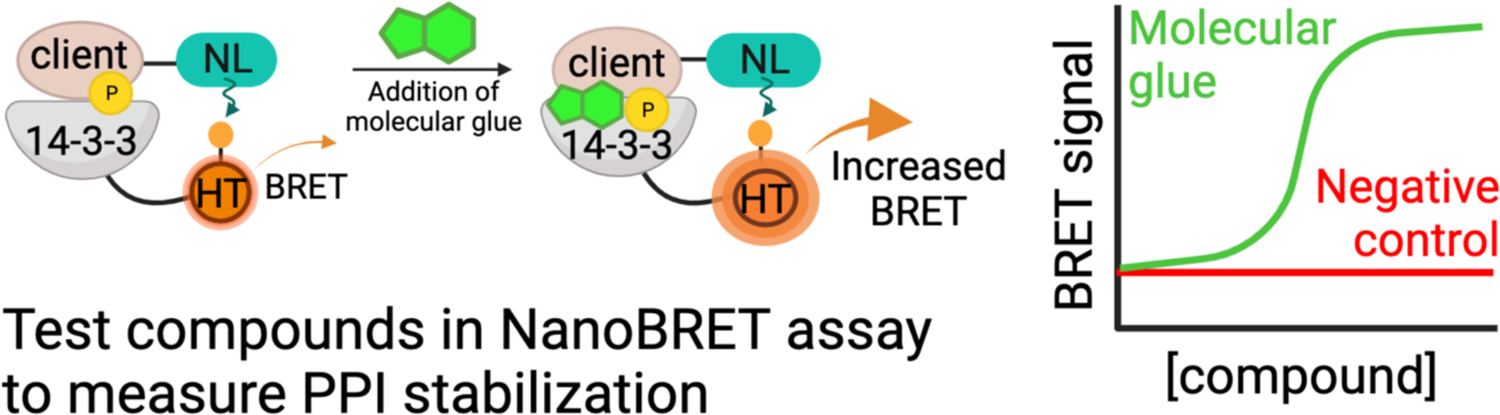

## 1. Introduction

### 1.1 Protein-Protein Interactions

Protein-protein interactions (PPIs) are crucial for various cellular processes and modulation of these interactions, by inhibition or stabilization, results in profound effects on cellular function.^1–4^ Molecular glues are small molecules that stabilize the interaction between two proteins by binding cooperatively to the PPI interface.^5–7^ Recent work on PPI stabilization has focused on developing molecular glues and bifunctional small molecules, called PROTACs, that link a ubiquitin ligase to a protein of interest to induce the protein’s degradation.^8,9^ Generally, PROTACs and molecular-glue degraders induce neomorphic interactions to degrade target proteins selectively.^6,7,9^ However, an intriguing and relatively underexplored approach lies in harnessing the potential of molecular glues as non-degradative stabilizers of PPIs.^5,10^ This novel perspective offers a promising approach to precisely modulate and stabilize PPIs for therapeutic interventions and for a deeper understanding of cellular processes.

Systematic technologies geared towards the identification of native PPI stabilizers^11–13^ tend to be target-directed, compared to cell-based technologies for molecular glue degraders.^14–16^ For instance, we have established biochemical fragment-based assays for the identification, validation, and optimization of PPI stabilizers. Our workflow employs a mass spectrometry (MS)-based site-directed disulfide tethering approach that has identified highly cooperative stabilizers for multiple 14-3-3/client complexes.^11,12,17^ During chemical optimization, PPI stabilizers are evaluated using MS- and fluorescence anisotropy (FA)-based assays. These technologies measure stabilization of 14-3-3 and a peptide derived from the client protein, facilitating screening and x-ray crystallography.^11,12,18^ With effective PPI stabilizers in hand, we sought a systematic approach to measure stabilization of full-length protein complexes. Ideally, such an approach would be suitable to lysate and cellular contexts. Here we describe the validation of NanoBRET assays to evaluate PPI stabilizers for the hub protein 14-3-3 bound to diverse protein partners.

### 1.2 The Hub Protein 14-3-3

14-3-3 modulates the function of hundreds of disease-relevant ‘client’ proteins through binding to linear sequences containing phosphorylated serine and threonine residues. Within the 14-3-3 family there are seven isoforms, each encoded by different genes.^19,20^ Notably, these isoforms exhibit pronounced sequence similarity within the phospho-binding groove, underscoring a shared mechanism of interaction with phosphorylated regions of client proteins.^21,22^ Peptides derived from the client proteins interact with the 14-3-3 binding groove via distinct conformations,^20,22,23^ including: (1) turning out of the groove after a phosphoserine/phosphothreonine (pS/pT) residue, typically guided by a proline in the +2 position, (2) binding straight through the phospho-binding groove, and (3) using a truncated sequence with a pS/pT at the penultimate residue of the protein. The three clients we describe here represent these three binding modes, with CRAF representing mode 1, TAZ reflecting mode 2, and Estrogen Receptor α (ERα) binding as mode 3 (Figure 1A; Supplementary Figure 2).^24–26^ Figure 1B illustrates the 14-3-3σ/client pocket we aim to target and example molecular glues.

**Figure 1:**
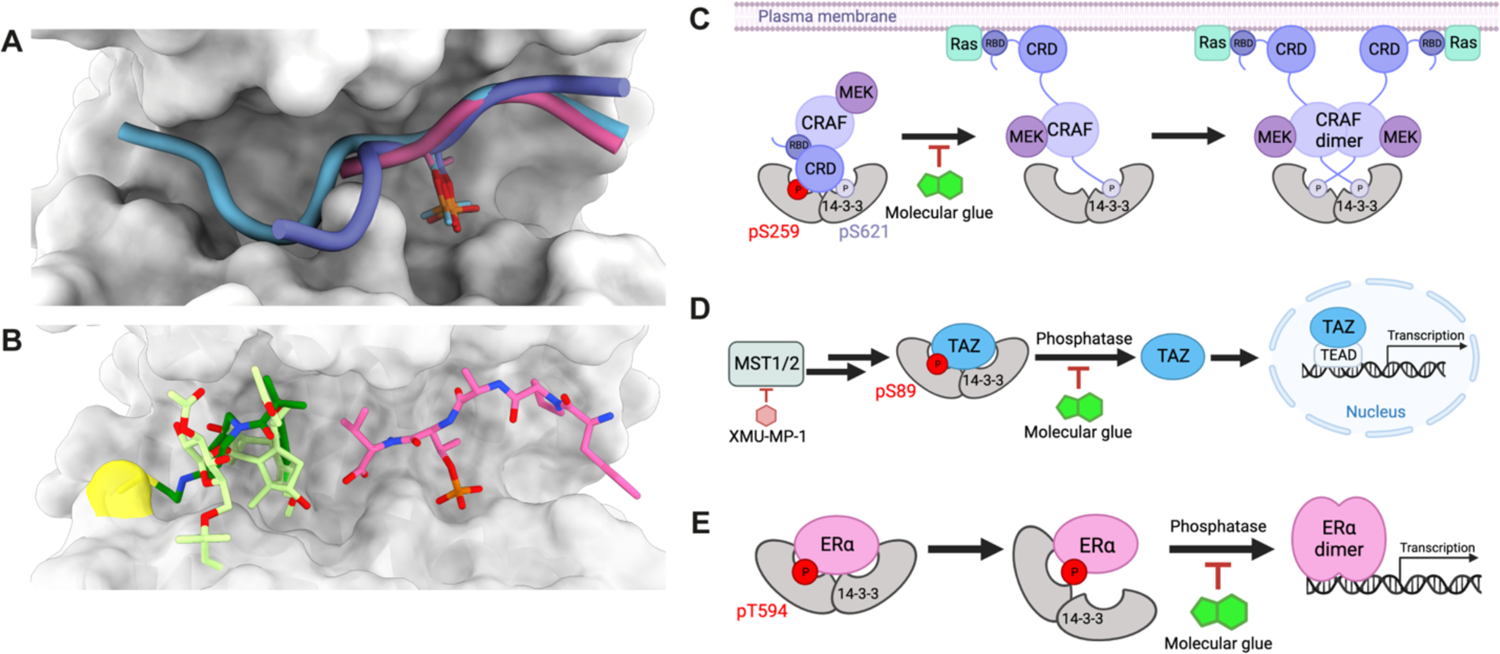
14-3-3σ/client binding and regulatory mechanisms. **A)** Crystal structure overlay of 14-3-3σ (grey) bound to CRAF (purple), ERα (pink), and TAZ (blue) peptides illustrating the three different binding modes. pS/T residues are shown to illustrate the overlap in this position of the peptide. PDBs: CRAF 4IHL, ERα 4JDD, TAZ 3MHR. **B)** Visualization of the molecular glue binding pocket within the 14-3-3σ phospho-binding groove. Both the fragment stabilizer (dark green) and Fusicoccin-A (light green) bind in the 14-3-3σ phospho-binding groove proximal to the ERα peptide (pink). 14-3-3σ C38 is highlighted in yellow. PDB 8AFN with Fusicoccin-A overlaid from PDB 4JDD. **C)** Proposed 14-3-3/CRAF mechanism. 14-3-3 binding to CRAF pS259 (red) maintains the inhibited state of CRAF. 14-3-3σ/CRAF pS259 molecular glues would prevent the opening and subsequent activation of CRAF. **D)** 14-3-3 regulation of TAZ in the Hippo signaling pathway. MST1/2 activity leads to the phosphorylation of TAZ at pS89 (red). In the nucleus, TAZ binds TEAD transcription factors and activates transcription. 14-3-3σ/TAZ pS89 molecular glues would localize TAZ in the cytoplasm, impeding its role as a transcription factor. **E)** 14-3-3 regulation of ERα. 14-3-3 binds ERα at pT594 (red) to prevent ERα dimerization and binding to DNA. 14-3-3σ/ERα pT594 molecular glues would prevent ERα dissociation and activation.

14-3-3 plays important biological roles in the regulation of CRAF, TAZ, and ERα. 14-3-3 is involved in both the inhibition and activation of CRAF, with CRAF pS259 acting as a regulatory switch that is only phosphorylated in the inhibited state. In the activation of CRAF, pS259 releases from 14-3-3 and can be dephosphorylated.^27–32^ Stabilizing the 14-3-3σ/CRAF pS259 interaction with molecular glues could therefore represent a novel method of inhibiting the MAPK pathway in various cancers (Figure 1C).^32^ The interaction between 14-3-3 and TAZ is a regulatory mechanism in the Hippo pathway. Phosphorylation of TAZ at S89 promotes association with 14-3-3, preventing TAZ translocation into the nucleus and activation as a transcription factor.^25,33^ Stabilizing the 14-3-3σ/TAZ pS89 interaction with molecular glues would inhibit the oncogenic role of TAZ in driving cell proliferation and survival (Figure 1D). 14-3-3 binds to ERα at the penultimate residue pT594. Binding to 14-3-3 inhibits ERα by preventing its dimerization and DNA binding. This interaction is crucial in the context of ER-positive breast cancer, where stabilizing the 14-3-3σ/ERα pT594 interaction may offer a potential therapeutic strategy to disrupt estrogen-driven tumorigenesis (Figure 1E).^26^ Not only do these clients represent different binding modes, but also different subcellular locations and functions.

Distinct classes of molecular-glue stabilizers of 14-3-3/client complexes have been discovered. Fusicoccane natural products and semisynthetic analogs,^20,26,34–36^ non-covalent synthetic compounds,^37,38^ and lysine-reactive fragments bind in the 14-3-3 phospho-binding groove and stabilize clients to all 14-3-3 isoforms.^39,40^ We have also identified disulfide fragments that bind to a cysteine residue at the periphery of the binding groove of the sigma isoform only and stabilize diverse 14-3-3α/client complexes.^11,12^ Structure-guided optimization towards 14-3-3/ERα stabilizers yielded cysteine-reactive molecules with EC_50_ values in the low μM range in biochemical assays and the ability to stabilize the complex by 120-fold, eg, K_D,app_ = 2 μM to 18 nM (Supplementary Figure 1).^18^ We and our colleagues have also developed 14-3-3α-specific stabilizers for CRAF and TAZ phosphopeptides. These three chemical series require cell-based validation of their ability to act as molecular glues for 14-3-3α and full-length proteins.

### 1.3 NanoBRET Assay

NanoLuciferase bioluminescence resonance energy transfer (NanoBRET) assays are widely adopted methods for measuring PPI and protein/small-molecule interactions in live cells.^30,41–44^ This innovative approach utilizes fusion proteins to quantify energy transfer, a process contingent upon the proximity between the tagged proteins of interest. NanoBRET assays have been effectively harnessed to measure PPI inhibition in cells and to measure formation of ubiquitin ligase/target complexes for PROTACs, showcasing their utility in probing the effects of modulating PPIs.^41,45,46^ However, to our knowledge, NanoBRET systems have not been used to measure PPI stabilization in the presence of molecular glues. We selected NanoBRET over related technologies such as cell-based fluorescence resonance energy transfer (FRET) or NanoLuc Binary Technology (NanoBiT) due to its improved sensitivity and geometric flexibility, respectively.^42,45,47,48^ Here we report a 384-well plate-based cellular NanoBRET method for the evaluation of 14-3-3σ/client molecular glues. The optimized assay allows for a statistically robust (Z’ = 0.7-0.95) increase in the BRET signal of 14-3-3σ/client interactions in the presence of small-molecule stabilizers. This 14-3-3σ/client stabilization NanoBRET methodology can be universally applied to characterize stabilizers for other PPIs, employing a similarly efficient and systematic approach.

## 2. Materials & Methods

### 2.1 Construct design and cloning

14-3-3σ and client proteins CRAF, TAZ, and ERα were cloned into the Promega NanoBRET vectors (Promega N1811) using Gibson Assembly (NEB E2611) with a GSSG linker between the protein and the tag. 17 residues were deleted from the C-terminus of 14-3-3σ to reduce dynamics of C-terminal tags; therefore, all 14-3-3σ constructs contain residues 1-231. NES-TAZ and NES-ERα constructs contain an engineered nuclear export signal (NES) on the N-terminus with a GSSG linker before the NanoLuc tag.

### 2.2 Compounds

CRAF-01-05 were synthesized in house and will be disclosed in due course. TAZ-01-03 were identified in a high-throughput screening campaign; the structures have not been disclosed (Ambagon Therapeutics). ERα-01 and ERα-02 were published as compounds 85 and 181, respectively.^18^ Fusicoccin-A was purchased from Enzo Life Sciences (BML-EI334-0001) and XMU-MP-1 was purchased from MedChemExpress (HY-100526).

### 2.3 Cell lines

HEK293T cells were purchased from ATCC (CRL-3216) and used as recommended. To avoid signal inhibition by endogenous, unlabeled proteins, we selected HEK293T cells because they express minimal to no 14-3-3σ, TAZ, and ERα. HEK293T cells also have an inactive MAPK pathway, suggesting that transfected CRAF would be phosphorylated at S259.

### 2.4 NanoBRET

NanoBRET assays were performed as described by Promega. Four constructs (two for ERα) were generated for each client protein to test in combination with four constructs of 14-3-3σ. Combination tests were performed at a 1:10 ratio of NanoLuc:HaloTag; ratios of 1:1, 1:10, 1:100, and 1:1000 were then tested with selected combinations. Plasmids were transfected using jetOPTIMUS transfection reagent following the manufacturer’s protocol and cells were plated in white, flat bottom TC-treated 384-well microplates (Corning 3570) after transfection in Gibco FluoroBrite DMEM with 4% fetal bovine serum (FBS; Sigma-Aldrich). ERα experiments were conducted with 4% charcoal dextran stripped FBS (Corning 35-072-CF). Each experiment was performed in triplicate and included no-acceptor controls and samples with the HaloTag NanoBRET 618 Ligand. Plates were read on an EnVision XCite 2105 plate reader at 618 nm (HaloTag) and 460nm (NanoLuc) (Filters: M647 CWL=647nm BW=75nm; M460 CWL=460nm BW=80nm Mirror: LUM/D585).

### 2.5 Fluorescence Anisotropy Measurements

ERα FA measurements were performed as previously described.^18^ For CRAF and TAZ, peptides were purchased from Elim Biopharmaceuticals, Inc (CRAF: QRSTpSTPNVH; TAZ: RSHpSSPASLQ). For CRAF, fluorescein-labeled peptides (5-FAM), full-length 14-3-3σ protein, the compounds (50 mM stock solution in DMSO) were diluted in buffer (10 mM HEPES, pH 7.5, 150 mM NaCl, 0.1% Tween20, 1 mg/mL Bovine Serum Albumin (BSA; Sigma-Aldrich). Final DMSO in the assay was always 1%. Dilution series of 14-3-3 proteins or compounds were made in black, flat-bottom 384-microwell plates (Greiner Bio-one 784900) in a final sample volume of 10 µL in triplicate. For compound titrations the initial 50 mM compound stock solutions in DMSO were diluted to 20mM in a 384-well master plate, followed by a serial 2-fold dilution. Then, 500 nl of the dilution series were transferred in the assay plates. A master mix containing 10 nM fluorescein-labeled peptide and 5 μM full-length 14-3-3σ (concentration at EC_20_ value of the protein-peptide complex) in buffer (10 mM HEPES, pH 7.5, 150 mM NaCl, 0.1% Tween20, 1 mg/mL BSA) was dispensed on the assay plates. The final volume per well was 10μL, with final top concentration of compounds dose response series at 1mM. Each compound was measured in triplicate, in two independent experiments. Fluorescence anisotropy measurements were performed directly and after overnight incubation at room temperature. Protein titrations were made by titrating 14-3-3σ in a 2-fold dilution series (starting at 250 µM) to a mix of fluorescein-labeled peptide (10 nM) and DMSO or compound (100 µM). Fluorescence anisotropy measurements were performed after overnight incubation at room temperature. Fluorescence anisotropy values were measured using a Molecular Devices ID5 plate reader (filter set λ_ex_: 485 ± 20 nm, λ_em_: 535 ± 25 nm; integration time: 50 ms; settle time: 0 ms; shake 5sec, medium, read height 3.00 mm, G factor = 1. Data reported are at endpoint. EC_50_ and apparent K_D_ values were obtained from fitting the data with a four-parameter logistic model (4PL) in GraphPad Prism. Data was obtained and averaged based on two independent experiments. TAZ FA experiments were done as described for CRAF with the following exceptions: final sample volume was 25 µL using 384-well plates (Corning 3575), final peptide concentration was 100 nM 5-FAM-TAZ, and final full-length 14-3-3σ protein concentration was 300 nM in the compound titration. Measurements were taken at 1 hour and 24 hours for compound titration and protein titration, respectively.

## 3. Procedure

We have developed a NanoBRET assay to test small molecule molecular glues of 14-3-3σ/client PPIs (Figure 2A). To test new PPIs, eight NanoLuc/HaloTag-fused protein constructs were cloned, and all eight combinations were tested. After selecting the desired combination, four ratios of NanoLuc:HaloTag plasmid were used to determine the optimal transfection ratio before testing compounds to measure their effects on PPI stabilization (Figure 2B).

**Figure 2:**
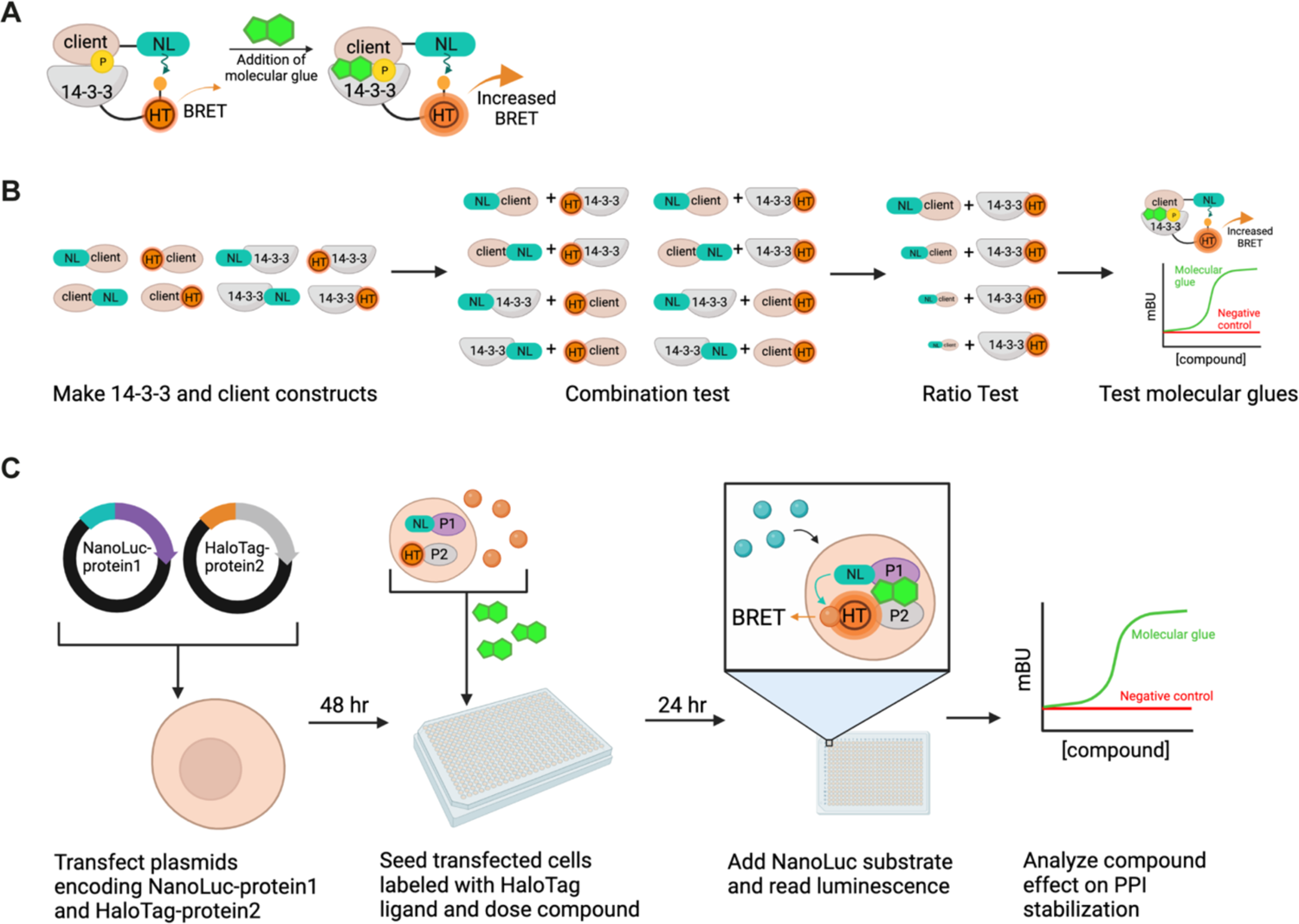
NanoBRET assay overview. **A)** Schematic representation of the 14-3-3/client NanoBRET assay, where one protein is tagged with NanoLuc (NL) and the other with HaloTag (HT). Addition of molecular glues is expected to increase the BRET signal. **B)** Steps to set up a NanoBRET assay for a new PPI. **C)** Procedure to set up NanoBRET assay.

### 3.1 Construct design

14-3-3σ and client proteins were cloned into NanoLuc and HaloTag vectors with N- or C-terminal tags. The constructs were named based on the placement of the tag, with “HaloTag-14-3-3σ” denoting N-terminal placement of the tag, and “14-3-3σ-HaloTag” denoting C-terminal placement. Because the C-terminus of ERα is crucial for binding to 14-3-3, ERα was only cloned with N-terminal tags. A nuclear export signal (NES; LPPLERLTL) was inserted at the N-terminus of TAZ and ERα for the “NES-client” constructs.^49^ The DNA binding domain (185-250) was truncated out of ERα to make the “ΔDBD-ERα” construct.

### 3.2 Assay Setup

Generally, we followed the manufacturer’s protocol.^41,50^ In short, plasmids were transfected in HEK293T cells in bulk for 48 hours. Ratios of NanoLuc:HaloTag plasmids were transfected by keeping the amount of HaloTag plasmid constant and decreasing the amount of NanoLuc plasmid. For example, a 1:10 transfection ratio could include 1 μg of HaloTag plasmid plus 0.1 μg of NanoLuc plasmid. The decrease in the amount of NanoLuc plasmid transfected is reflected in the amount of protein expressed as measured by western blot (Supplementary Figure 3A). Cells were trypsinized and counted before adjusting the cell density to 250,000 cells/mL and adding HaloTag ligand (1 μL ligand/1 mL of media). A no-HaloTag-ligand control was also made wherein DMSO was added instead of ligand at the same % v/v (1 μL/1 mL). 30 μL of the cell/HaloTag ligand or DMSO mixture was added to each well in a 384-well assay plate. Compound (or DMSO) was diluted in media to 4x the final concentration. Then, 10 μL of this 4x compound stock was added to corresponding wells. All samples had the same final percent DMSO of less than 0.5%. After 24 hours, NanoLuc substrate was added, and assay plates were read to measure luminescence in the NanoLuc and HaloTag channels (Figure 2C).

An alternative approach can be used to measure NanoBRET in lysate using the detergent digitonin. This approach can be used if the compounds are not cell permeable or are cytotoxic at the desired concentration range. Digitonin permeabilizes the outer membrane to release cytosolic proteins into the media (Supplementary Figure 3B). As exemplified for CRAF, we added digitonin at 200 μg/mL after seeding cells and before dosing with compound.

### 3.3 Data Analysis

The following formulas were used to calculate the raw milliBRET units (mBU), mean corrected mBU, and fold change. Data was then analyzed and visualized in GraphPad Prism to obtain EC_50_ values by fitting the data with a 4PL model.

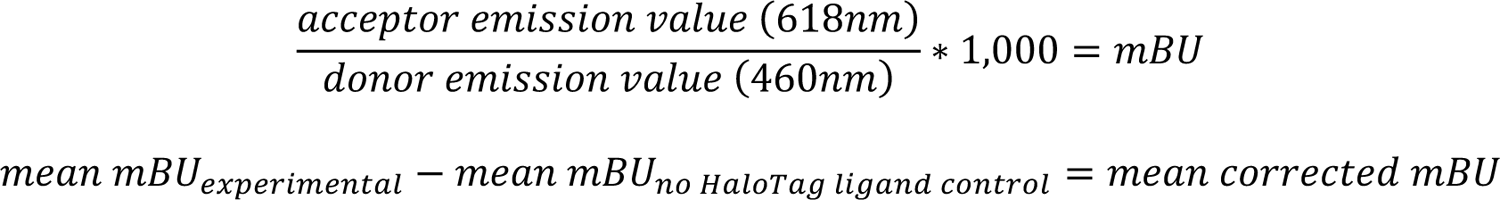

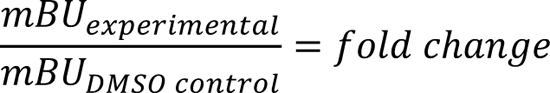

## 4. Results

### 4.1 CRAF/14-3-3σ NanoBRET

To measure cysteine-targeting stabilizers of the 14-3-3σ/CRAF pS259 interaction, eight constructs were cloned for 14-3-3σ/CRAF: HaloTag-14-3-3σ, 14-3-3σ-HaloTag, NanoLuc-14-3-3σ, 14-3-3σ-NanoLuc, HaloTag-CRAF, CRAF-HaloTag, NanoLuc-CRAF, and CRAF-NanoLuc. With these constructs, eight combinations were tested at a 1:10 NanoLuc:HaloTag transfection ratio (Figure 3A). Combinations where the NanoLuc was fused to 14-3-3σ and the HaloTag fused to CRAF resulted in low BRET signal. Two combinations, NanoLuc-CRAF/14-3-3σ-HaloTag and CRAF-NanoLuc/14-3-3σ-HaloTag (Z’ = 0.91 and 0.92, respectively), were selected for ratio testing. CRAF-NanoLuc/14-3-3σ-HaloTag gave higher signal and was chosen to move forward with. In these ratio tests, 1:10 and 1:100 CRAF-NanoLuc:14-3-3σ-HaloTag gave the highest signals (Z’ = 0.90, Z’ = 0.95, respectively; Figure 3B). Ratios of 1:1 and 1:1,000 were excluded due to low signal and high background, respectively.

**Figure 3:**
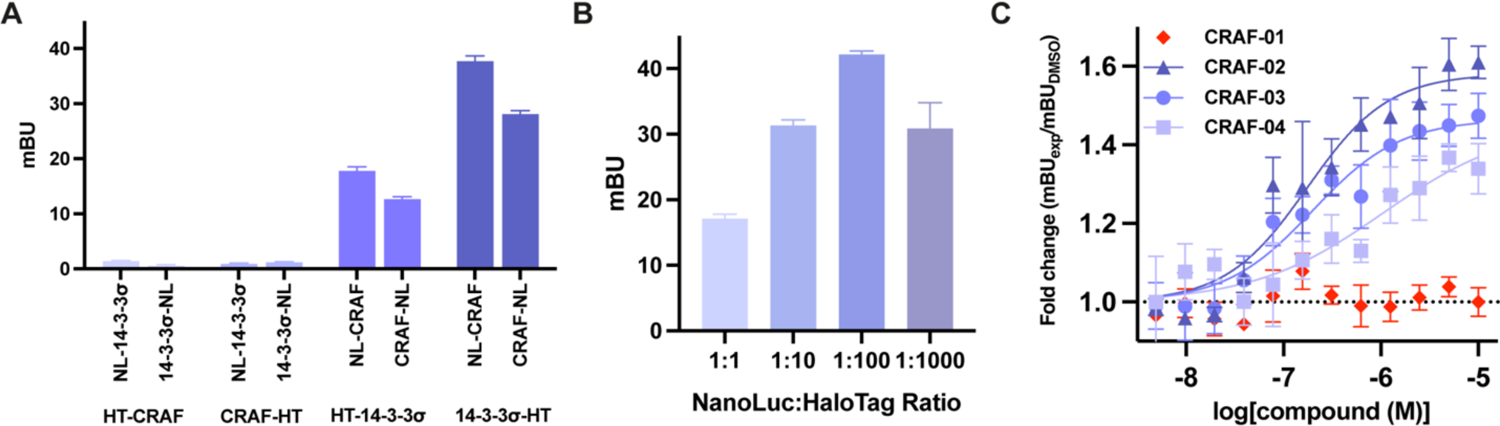
NanoBRET results for CRAF/14-3-3σ. **A)** Combination test (n = 3). 14-3-3σ with a C-terminal HaloTag and CRAF with an N- or C-terminal NanoLuc tag resulted in the highest signal. **B)** CRAF-NanoLuc/14-3-3σ-HaloTag ratio test (n = 3). **C)** CRAF-NanoLuc/14-3-3σ-HaloTag NanoBRET data with 4 compounds, three stabilizers (CRAF-02-04) and one negative control (CRAF-01) (n = 3). Data shown in C are representative of 5 biological replicates.

When using NanoBRET to identify molecular glues, careful selection of a combination and ratio is essential to ensure that the protein complex is not completely formed, such that it maintains sensitivity to stabilization by small molecules. Thus, we evaluated both 1:10 and 1:100 in the presence of stabilizers. 1:10 resulted in higher fold-stabilization with molecular glues than 1:100 (Supplementary Figure 4A); thus, we proceeded with this condition.

We tested three compounds, initially identified as molecular glues through MS and FA assays, along with one negative control, in the CRAF/14-3-3σ NanoBRET system. The negative control was structurally related to the stabilizers but inactive in the FA assay. The molecular glues exhibited up to 1.6-fold increase in NanoBRET signal, and EC_50_ values in the sub-μM to low-μΜ range (Figure 3C). The fold increase in signal and the Z’ values were similar between the NanoBRET and FA assays, suggesting similar sensitivity. These compounds were dependent on the presence of C38 in 14-3-3σ, as mutation of this residue rendered the compounds ineffective in stabilizing the interaction (Supplementary Figure 4B).

In comparing the EC_50_ values obtained from FA and NanoBRET, it is noteworthy that FA was performed in vitro with a fully phosphorylated CRAF pS259 peptide with the 14-3-3σ concentration kept at its EC_20_, while NanoBRET was performed in cells with the full-length CRAF that may not have been fully phosphorylated. Moreover, the degree of phosphorylation of CRAF may increase over the time of compound incubation, since increased 14-3-3 binding protects clients from dephosphorylation. These factors make it difficult to quantitatively compare EC_50_ values from the biochemical and cell-based binding assays. Nevertheless, the trends in EC_50_ values track between NanoBRET and FA, with the negative control CRAF-01 exhibiting a non-stabilizing effect. In contrast, CRAF-02 emerged as the most potent stabilizer with an EC_50_ value of 0.18 μM and the highest maximum signal. CRAF-03 had a similar EC_50_ value of 0.20 μM, but a lower maximum signal. CRAF-04 demonstrated a comparatively diminished stabilizing capacity with an EC_50_ value of 1.1 μM, making it the weakest of the three molecular glues (Table 1). The difference in signal maximum was reproducible and consistent with the FA data (Supplementary Figure 1A-E). The calculated EC_50_ values from the NanoBRET assay for CRAF-02 and CRAF-03 were lower than those calculated from FA assays (Table 1), but the rank order was maintained.

**Table 1:**
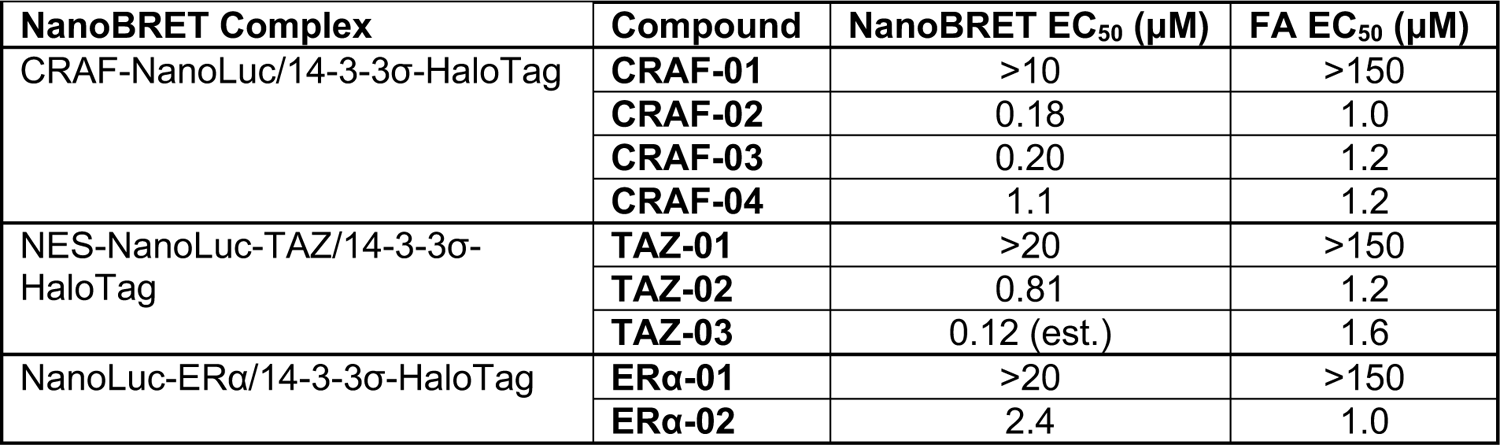
EC_50_ values for 14-3-3σ/client NanoBRET and FA (n = 3).

| NanoBRET Complex | Compound | NanoBRET EC<sub>50</sub> (μM) | FA EC<sub>50</sub> (μM) |
| --- | --- | --- | --- |
| CRAF-NanoLuc/14-3-3σ-HaloTag | **CRAF-01** | >10 | >150 |
|  | **CRAF-02** | 0.18 | 1.0 |
|  | **CRAF-03** | 0.20 | 1.2 |
|  | **CRAF-04** | 1.1 | 1.2 |
| NES-NanoLuc-TAZ/14-3-3σ-HaloTag | **TAZ-01** | >20 | >150 |
|  | **TAZ-02** | 0.81 | 1.2 |
|  | **TAZ-03** | 0.12 (est.) | 1.6 |
| NanoLuc-ERα/14-3-3σ-HaloTag | **ERα-01** | >20 | >150 |
|  | **ERα-02** | 2.4 | 1.0 |

Using the procedure described above, we introduced the detergent digitonin into the assay to permeabilize the cell membrane, releasing the NanoBRET pair into the media. In the cellular NanoBRET assay, CRAF stabilizers were tested up to 10 μΜ based on their cytotoxicity in HEK293T (data not shown). To compare the lysate and cell-based assays, we measured CRAF-05 activity in the CRAF/14-3-3σ NanoBRET with four doses, from 1 – 250 μM in lysate and 1 – 10 μM in cells. The results in both assays were similar, with CRAF-05 showing significant stabilization even at 1 μM. The lower signal ratio at the maximum concentrations (1.3-vs 1.5-fold) led us to use the cell-based assay going forward (Supplementary Figure 4C). Thus, PPI stabilization can be measured either in cells or in lysates using the same transfection ratio and assay formats.

### 4.2 TAZ/14-3-3σ NanoBRET

The following four constructs were cloned to test both donor and acceptor tags on the N- or C-terminus of TAZ: HaloTag-TAZ, TAZ-HaloTag, NanoLuc-TAZ, and TAZ-NanoLuc. To aid in developing the 14-3-3σ/TAZ NanoBRET system, we used an MST1/2 inhibitor, XMU-MP-1 (“XMU”), at 60 μM to prevent phosphorylation of TAZ, which in turn inhibited the 14-3-3σ/TAZ interaction (Figure 1D). After testing all eight NanoBRET combinations of 14-3-3σ and TAZ, we chose to move forward with NanoLuc-TAZ/14-3-3σ-HaloTag because it produced the largest BRET signal (24 mBU) with the most significant difference between untreated and XMU treated samples (fold change = 1.4, Z’ = 0.84; Figure 4A). NanoLuc:HaloTag ratios were then evaluated, and 1:10 gave the largest fold-change between untreated and XMU treated samples (fold-change = 1.4, Z’ = 0.90; Figure 4B).

**Figure 4:**
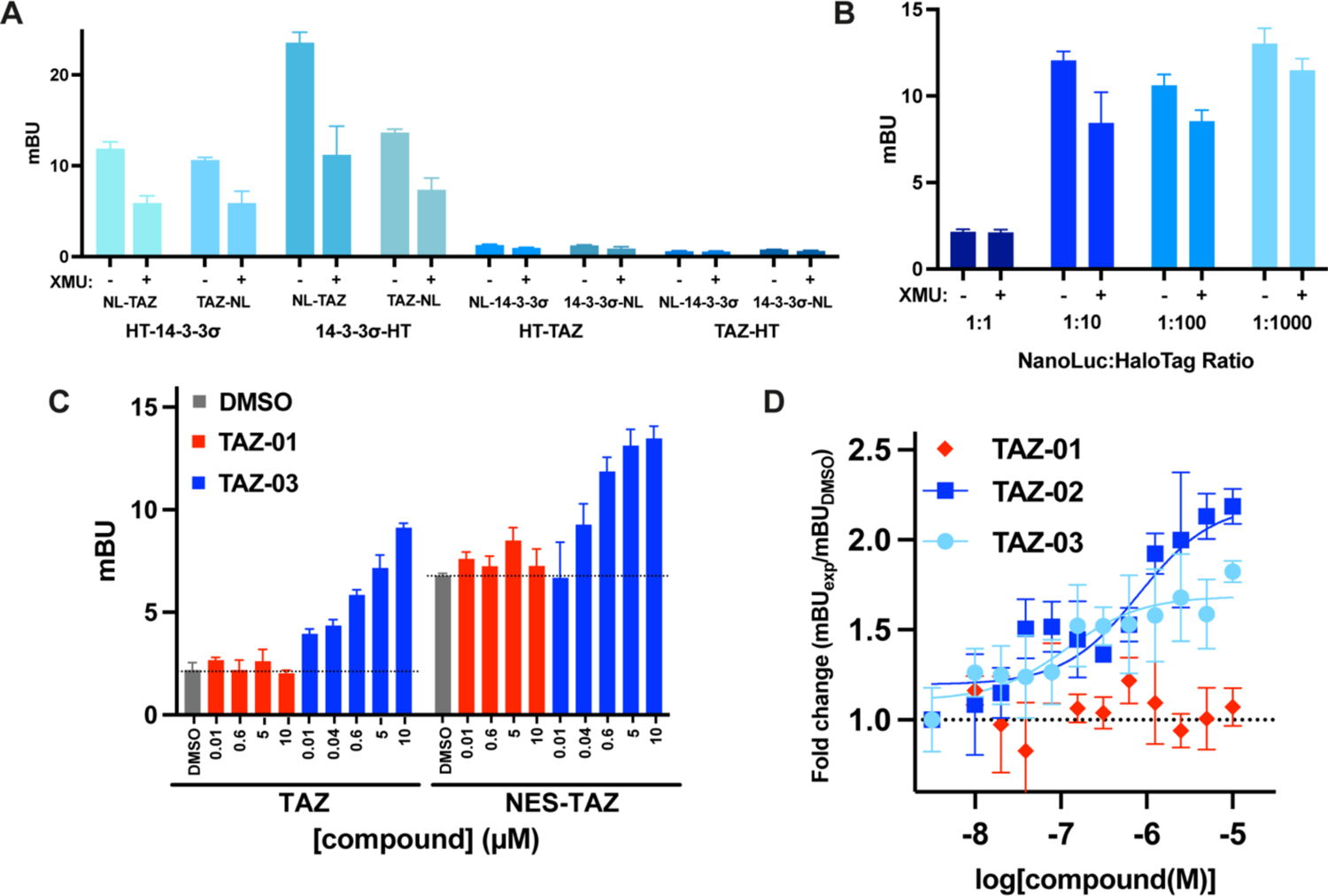
NanoBRET results for TAZ/14-3-3σ. **A)** Combination test comparing untreated and XMU (60 μM) treated samples (n = 3). NanoLuc-TAZ/14-3-3σ-HaloTag gave the highest signal and largest fold change with XMU. **B)** NanoLuc-TAZ/14-3-3σ-HaloTag ratio test (n = 3). **C)** Comparison of TAZ and NES-TAZ constructs with one inactive compound (TAZ-01) and one stabilizer (TAZ-02) (n = 3). A subset of tested compound concentrations is shown. **D)** NES-TAZ/14-3-3σ NanoBRET results with two TAZ/14-3-3σ stabilizers (TAZ-02 and TAZ-03) and TAZ-01 with all data points in the dilution series shown (n = 3). Data shown in C and D are representative of 3 biological replicates.

14-3-3σ controls trafficking of TAZ between the cytoplasm and the nucleus, so we evaluated whether manipulating NanoLuc-TAZ subcellular localization would alter the BRET signal. We engineered a nuclear export signal (NES) into the N-terminus of the TAZ construct before the NanoLuc tag. With this NES-NanoLuc-TAZ (NES-TAZ) construct, the basal NanoBRET signal increased 3.1-fold (Figure 4C). While the raw mBU signal varied between biological replicates, perhaps due to transfection efficiency, the BRET signal was consistently larger with the NES construct.

We compared the 14-3-3σ/TAZ complex formation with C38-reactive molecular glues, TAZ-02 and TAZ-03, along with an inactive compound, TAZ-01 (Supplementary Figure 5). The NES-TAZ construct led to increased mBU values but decreased the fold-change compared to NanoLuc-TAZ itself. For instance, TAZ-02 increased the BRET signal by 4.2-fold and 2.2-fold with NanoLuc-TAZ and NES-TAZ, respectively (Figure 4C). Nevertheless, the compound dose-responses yielded similar EC_50_ values with both TAZ constructs. The negative control TAZ-01 had no effect on the BRET signal, while TAZ-02 had an EC_50_ of 810 nM; TAZ-03 did not reach a plateau, with an estimated EC_50_ of 120 nM (Figure 4D). In FA assays, TAZ-02 and TAZ-03 performed similarly to each other with EC_50_ values of 1.2 μM and 1.6 μM, respectively (Supplementary Figure 1F-I; Table 1). Thus, the TAZ/14-3-3σ NanoBRET assays confirmed the stabilizing effects of these molecular glues.

### 4.3 ERα/14-3-3σ NanoBRET

ERα binds 14-3-3σ at its penultimate residue, pT594. Therefore, only N-terminal tag constructs were developed for ERα to avoid interference with 14-3-3σ binding. ERα combination and ratio tests were performed in the presence of a natural product stabilizer, Fusicoccin-A (FC-A) at 30 μΜ. Similar to CRAF and TAZ, little to no signal was measured for the combinations in which 14-3-3σ was tagged with NanoLuc. The NanoLuc-ERα plus 14-3-3σ-HaloTag combination gave the largest signal and fold stabilization for the positive control FC-A (Z’ = 0.79, Figure 5A). In the ratio test, 1:10 was the only combination where there was a measurable difference between DMSO and FC-A (Z’ = 0.96, Figure 5B). Therefore, this ratio was selected to advance to the subsequent testing phase for new 14-3-3σ specific, cysteine-reactive molecular glues (Supplementary Figure 6A). The most potent molecular glue, ERα-02, resulted in 2.5-fold stabilization with an EC_50_ of 2.4 μM (Figure 5C). In FA dose-responses, ERα-02 had a similar EC_50_ value of 1.0 μM when tested with the ERα peptide (Supplementary Figure 1J,K).^18^ The inactive compound, ERα-01, had no effect in NanoBRET or FA dose response assays (Figure 5C, Table 1). FC-A resulted in low fold stabilization (Figure 5A,B) likely because binding of the NanoBRET pair was diluted by binding to other 14-3-3 isoforms and clients. Because HEK293T cells did not contain endogenous 14-3-3σ, the sigma-specific stabilizer, ERα-02, was highly effective at binding to the transfected 14-3-3σ.

**Figure 5:**
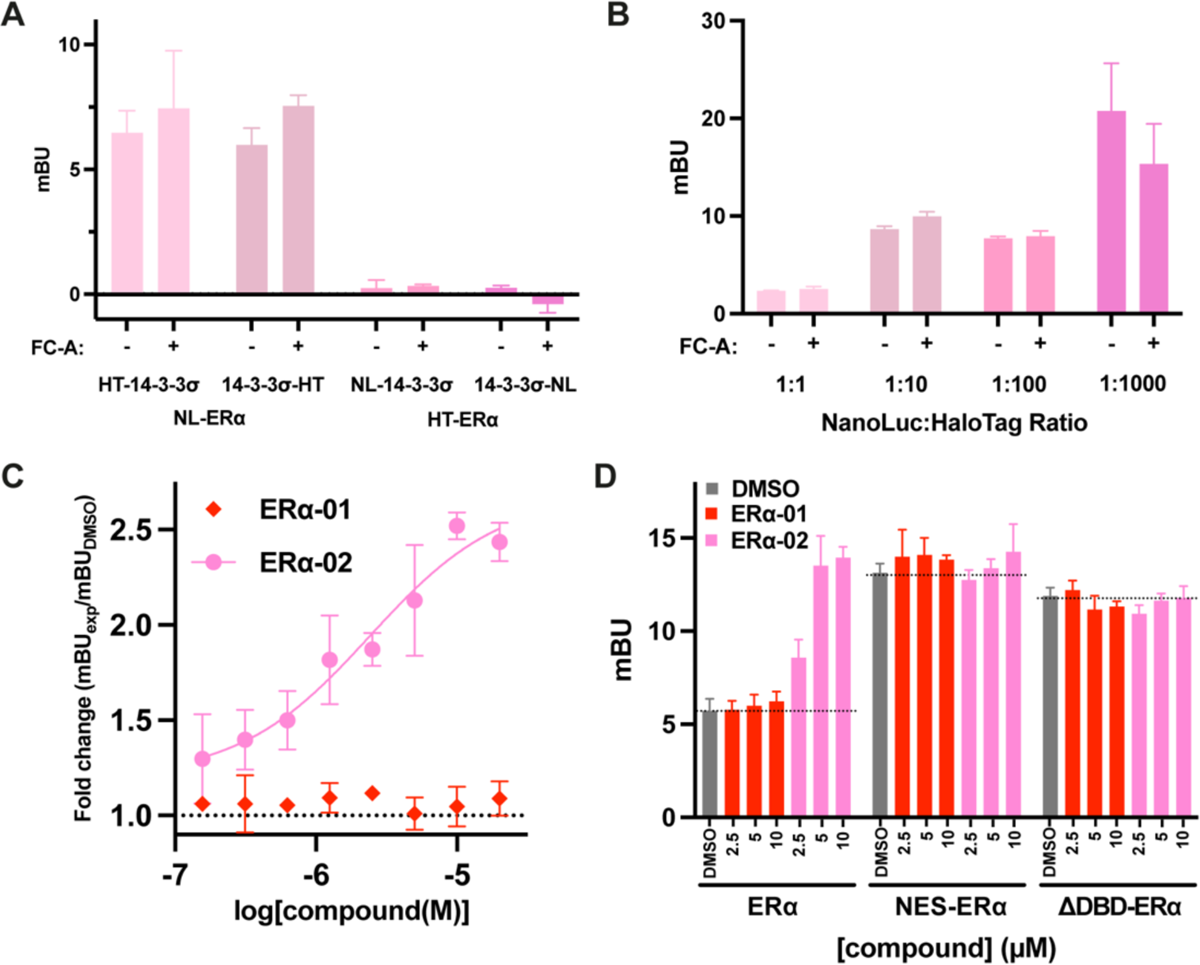
NanoBRET results for ERα/14-3-3σ. **A)** Combination test comparing untreated and FC-A (30 μΜ) treated samples (n = 3). NanoLuc-ERα/14-3-3σ-HaloTag gave the only significant difference with DMSO vs FC-A treatment. **B)** NanoLuc-ERα/14-3-3σ-HaloTag ratio test (n = 3). **C)** ERα/14-3-3σ NanoBRET results with one molecular-glue stabilizer (ERα-02) and one inactive compound (ERα-01) (n = 3). **D)** Comparison of different ERα constructs with ERα-01 and ERα-02 (n = 3). Data shown in C and D are representative of 3 biological replicates.

Since ERα is also a transcription factor, a significant fraction could be localized in the nucleus, resulting in a low NanoBRET signal. We therefore engineered two cytoplasmic constructs containing an NES at the N-terminus (“NES-ERα”) or removing the DNA-binding domain (aa 185-250; “ΔDBD-ERα”). While both methods increased the BRET signal, from 5-10 to 10-15 mBU, the stabilizer dose response effects were diminished (Figure 5D). Although these methods of increasing the NanoBRET signal did not improve the 14-3-3σ/ERα NanoBRET assay for ERα-02, they are useful techniques to apply when troubleshooting low NanoBRET signal for other PPI NanoBRET systems, as seen with TAZ.

A benefit of developing the ERα/14-3-3σ NanoBRET assay in HEK293T cells was the low endogenous expression of both proteins. We extended these experiments to an ER-positive breast cancer cell line, MCF7, where both ERα and 14-3-3σ were expressed, and MDA-MB-231, an ER-negative breast cancer cell line, where only 14-3-3σ was endogenously expressed. Comparing the ERα/14-3-3σ NanoBRET assay in these three cell lines, HEK293T cells had the largest signal window with minimal background (Supplementary Figure 6B-D). Thus, using a cell line that does not endogenously express the proteins of interest ensured that the tagged proteins were not outcompeted by endogenous proteins.

## 5. Discussion

### 5.1 14-3-3σ NanoBRET Summary and Workflow

In developing analogous NanoBRET assays for three 14-3-3σ/client complexes, we learned that the HaloTag acceptor was always preferred on 14-3-3σ and that varying the subcellular localization of the client protein sometimes improved the BRET signal, depending on the biology of the complex (Table 2). Traditional NanoBRET assay development conventionally involves a four-step workflow: (1) generating eight constructs of 14-3-3σ and the client tagged with NanoLuc or HaloTag, (2) testing eight combinations, (3) evaluating four ratios of the best combination, and (4) subsequently testing molecular glues (Figure 2B). Leveraging the insights gained from our results across 14-3-3σ/client systems, we propose a streamlined workflow that condenses the NanoBRET assay design for 14-3-3σ/client interactions into three efficient steps:

1. constructing two client constructs (or one depending on client binding restrictions, e.g. ERα),
2. test two combinations, and (3) evaluate molecular glues (Figure 6). This simplified workflow promises expedited discovery of 14-3-3σ/client molecular-glue stabilizers, offering a potentially impactful resource for diverse therapeutic applications.

**Figure 6:**
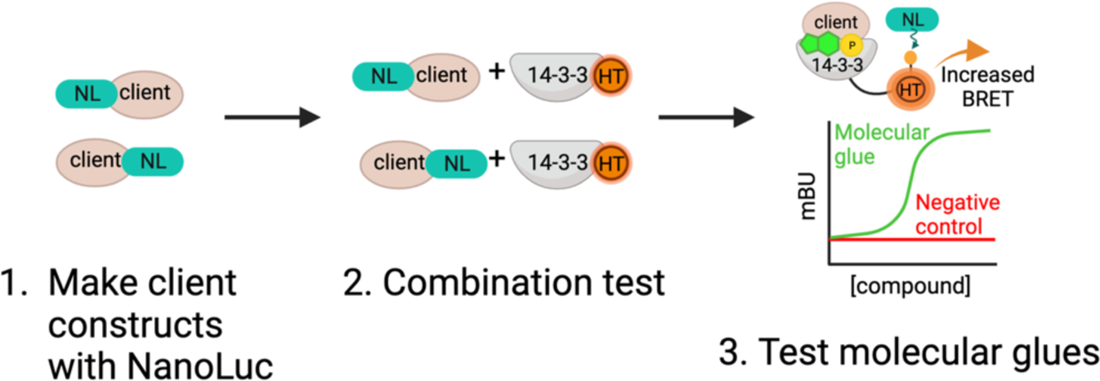
Simplified NanoBRET workflow for identification of 14-3-3σ/client molecular glues.

**Table 2:** Tag placement for 14-3-3σ/CRAF, TAZ, and ERα NanoBRET assays.

| 14-3-3σ client | NanoLuc | HaloTag |
| --- | --- | --- |
| CRAF | CRAF (C-terminus) | 14-3-3σ (C-terminus) |
| TAZ | TAZ (N-terminus) | 14-3-3σ (C-terminus) |
| ERα | ERα (N-terminus) | 14-3-3σ (C-terminus) |

### 5.2 Assay Throughput

NanoBRET assays to discover and characterize molecular glues can be set up in a low-, medium-, or high-throughput format depending on materials and instruments employed. Generally, we have run approximately 200-2,000 samples at a time. Because optimizing the transfection ratio is necessary to detect PPI stabilizers, the NanoBRET assay requires a transient transfection. This manual step reduces throughput and precision; in particular, raw mBU values can vary between experiments. Nevertheless, with automation for cell seeding and compound dosing, the NanoBRET assay for PPI stabilizers can be performed in a high-throughput manner.

### 5.3 Conclusions

We describe the development of a NanoBRET assay designed to characterize molecular-glue stabilizers of 14-3-3σ/client PPIs. Building from the published NanoBRET protocol,^50^ we have identified multiple factors that required careful evaluation. Specific to PPI stabilizers, it is critical to select a combination of protein constructs and transfection ratio that leads to a measurable but incomplete formation of the PPI, such that the assay can detect an increase in BRET signal. Additional general considerations include optimizing the cell permeability and colocalization of the protein partners. For example, digitonin releases cytoplasmic proteins into the media, allowing for higher compound concentrations and effectively transforming the assay into a lysate assay. Furthermore, adding a nuclear export signal increased mBU values for the transcription factor/14-3-3σ complexes, suggesting that localization motifs should be considered if the BRET signal is low. Lastly, choice of cell line should account for the endogenous expression levels of the proteins of interest. With these considerations, our NanoBRET results closely mirror the sensitivity of FA and the NanoBRET EC_50_ values were consistent with the FA EC_50_ values measured for diverse molecular glues. Thus, NanoBRET is a valuable tool to characterize stabilizers targeting 14-3-3σ/client interactions. Notably, the distinct advantage of NanoBRET is the physiological relevance of the assay, as it is performed in a cellular environment with full-length proteins.

This comprehensive method can be efficiently adapted and streamlined for the exploration of other 14-3-3σ client proteins. Moreover, the systematic approach can be readily applied to other hub protein systems and to discover novel molecular glues. As research in this domain progresses, the integration of NanoBRET assays for PPI stabilization could significantly broaden the repertoire of techniques available for studying PPIs and their implications in cellular processes.

## Abbreviations

PPI: Protein-protein interaction

PROTAC: Proteolysis targeting chimera

MS: Mass spectrometry

FA: Fluorescence anisotropy

pS/T: Phosphoserine/phosphothreonine

NanoBRET: NanoLuciferase bioluminescence resonance energy transfer

FRET: Fluorescence resonance energy transfer

NanoBiT: NanoLuc binary technology

4PL: 4 parameter logistic

NES: Nuclear export signal

mBU: milliBRET units

XMU: XMU-MP-1

FC-A: Fusicoccin-A

ΔDBD-ERα: ERα without DNA binding domain

## Author Contributions

H.R.V. and J.M.V. contributed equally to this work. The manuscript was written through contributions of all authors. All authors have given approval to the final version of the manuscript.

## Funding

This research was funded by the National Institutes of Health grant GM147696.

## Declaration of Conflicting Interests

The authors declare the following competing financial interest: M.R.A. is a co-founder and director of Ambagon Therapeutics.

## Acknowledgements

The authors would like to thank Ambagon Therapeutics for providing 14-3-3σ/TAZ stabilizers. We also thank Dr. Luc Brunsveld and Dr. Christian Ottmann for collaborative discussion. We would like to thank Dr. Ziwen Jiang for assay support, Dr. Andrew J Ambrose for useful discussion, and Dr. Jezrael Lafuente Revalde and Zain Alam for instrument training. We also thank Sean Woods for assay support and discussion.

## References

1. Arkin, M. R., Tang, Y. & Wells, J. A. Small-Molecule Inhibitors of Protein-Protein Interactions: Progressing toward the Reality. Chem. Biol. 21, 1102–1114 (2014).

2. Ran, X. & Gestwicki, J. E. Inhibitors of Protein-Protein Interactions (PPIs): An Analysis of Scaffold Choices and Buried Surface Area. Curr. Opin. Chem. Biol. 44, 75–86 (2018).

3. Andrei, S. A. et al. Stabilization of protein-protein interactions in drug discovery. Expert Opin. Drug Discov. 12, 925–940 (2017).

4. Garlick, J. M. & Mapp, A. K. Selective Modulation of Dynamic Protein Complexes. Cell Chem. Biol. 27, 986–997 (2020).

5. Schreiber, S. L. The Rise of Molecular Glues. Cell 184, 3–9 (2021).

6. Dong, G., Ding, Y., He, S. & Sheng, C. Molecular Glues for Targeted Protein Degradation: From Serendipity to Rational Discovery. J. Med. Chem. 64, 10606–10620 (2021).

7. Sasso, J. M. et al. Molecular Glues: The Adhesive Connecting Targeted Protein Degradation to the Clinic. Biochemistry 62, 601–623 (2023).

8. Békés, M., Langley, D. R. & Crews, C. M. PROTAC targeted protein degraders: the past is prologue. Nat. Rev. Drug Discov. 21, 181–200 (2022).

9. Chamberlain, P. P. & Hamann, L. G. Development of targeted protein degradation therapeutics. Nat. Chem. Biol. 15, 937–944 (2019).

10. Soini, L., Leysen, S., Davis, J. & Ottmann, C. Molecular glues to stabilise protein-protein interactions. Curr. Opin. Chem. Biol. 69, 102169 (2022).

11. Kenanova, D. N. et al. A Systematic Approach to the Discovery of Protein–Protein Interaction Stabilizers. ACS Cent. Sci. 9, 937–946 (2023).

12. Sijbesma, E. et al. Site-Directed Fragment-Based Screening for the Discovery of Protein– Protein Interaction Stabilizers. J. Am. Chem. Soc. 141, 3524–3531 (2019).

13. Sijbesma, E. et al. Structure-based evolution of a promiscuous inhibitor to a selective stabilizer of protein–protein interactions. Nat. Commun. 11, 3954 (2020).

14. Toriki, E. S. et al. Rational Chemical Design of Molecular Glue Degraders. ACS Cent. Sci. 9, 915–926 (2023).

15. King, E. A. et al. Chemoproteomics-enabled discovery of a covalent molecular glue degrader targeting NF-κB. Cell Chem. Biol. 30, 394–402.e9 (2023).

16. Mayor-Ruiz, C. et al. Rational discovery of molecular glue degraders via scalable chemical profiling. Nat. Chem. Biol. 16, 1199–1207 (2020).

17. Hallenbeck, K. K. et al. A Liquid Chromatography/Mass Spectrometry Method for Screening Disulfide Tethering Fragments. SLAS Discov. Adv. Sci. Drug Discov. 23, 183–192 (2018).

18. Konstantinidou, M. et al. Structure-Based Optimization of Covalent, Small-Molecule Stabilizers of the 14-3-3σ/ERα Protein–Protein Interaction from Nonselective Fragments. J. Am. Chem. Soc. 145, 20328–20343 (2023).

19. Obsilova, V. & Obsil, T. The 14-3-3 Proteins as Important Allosteric Regulators of Protein Kinases. Int. J. Mol. Sci. 21, 8824 (2020).

20. Stevers, L. M. et al. Modulators of 14-3-3 Protein–Protein Interactions. J. Med. Chem. 61, 3755–3778 (2018).

21. Obsil, T. & Obsilova, V. Structural basis of 14-3-3 protein functions. Semin. Cell Dev. Biol. 22, 663–672 (2011).

22. Pennington, K. L., Chan, T. Y., Torres, M. P. & Andersen, J. L. The dynamic and stress-adaptive signaling hub of 14-3-3: emerging mechanisms of regulation and context-dependent protein–protein interactions. Oncogene 37, 5587–5604 (2018).

23. Manschwetus, J. T. et al. Binding of the Human 14-3-3 Isoforms to Distinct Sites in the Leucine-Rich Repeat Kinase 2. Front. Neurosci. 14, (2020).

24. Molzan, M. et al. Impaired Binding of 14-3-3 to C-RAF in Noonan Syndrome Suggests New Approaches in Diseases with Increased Ras Signaling. Mol. Cell. Biol. 30, 4698–4711 (2010).

25. Sijbesma, E. et al. Identification of Two Secondary Ligand Binding Sites in 14-3-3 Proteins Using Fragment Screening. Biochemistry 56, 3972–3982 (2017).

26. De Vries-van Leeuwen, I. J., et al. Interaction of 14-3-3 proteins with the estrogen receptor alpha F domain provides a drug target interface. Proc. Natl. Acad. Sci. U. S. A. 110, 8894– 8899 (2013).

27. Lavoie, H. & Therrien, M. Regulation of RAF protein kinases in ERK signalling. Nat. Rev. Mol. Cell Biol. 16, 281–298 (2015).

28. Park, E. et al. Architecture of autoinhibited and active BRAF–MEK1–14-3-3 complexes. Nature 575, 545–550 (2019).

29. Kondo, Y. et al. Cryo-EM structure of a dimeric B-Raf:14-3-3 complex reveals asymmetry in the active sites of B-Raf kinases. Science 366, 109–115 (2019).

30. Martinez Fiesco, J. A., Durrant, D. E., Morrison, D. K. & Zhang, P. Structural insights into the BRAF monomer-to-dimer transition mediated by RAS binding. Nat. Commun. 13, 486 (2022).

31. Sun, Q. & Wang, W. Structures of BRAF–MEK1–14-3-3 sheds light on drug discovery. Signal Transduct. Target. Ther. 4, 1–2 (2019).

32. Park, E. et al. Cryo-EM structure of a RAS/RAF recruitment complex. Nat. Commun. 14, 4580 (2023).

33. Freeman, A. K. & Morrison, D. K. 14-3-3 Proteins: Diverse Functions in Cell Proliferation and Cancer Progression. Semin. Cell Dev. Biol. 22, 681–687 (2011).

34. Molzan, M. et al. Stabilization of Physical RAF/14-3-3 Interaction by Cotylenin A as Treatment Strategy for RAS Mutant Cancers. ACS Chem. Biol. 8, 1869–1875 (2013).

35. Ballone, A., Centorrino, F. & Ottmann, C. 14-3-3: A Case Study in PPI Modulation. Molecules 23, 1386 (2018).

36. Wolter, M. et al. Selectivity via Cooperativity: Preferential Stabilization of the p65/14-3-3 Interaction with Semisynthetic Natural Products. J. Am. Chem. Soc. 142, 11772–11783 (2020).

37. Richter, A., Rose, R., Hedberg, C., Waldmann, H. & Ottmann, C. An Optimised Small-Molecule Stabiliser of the 14-3-3–PMA2 Protein–Protein Interaction. Chem. – Eur. J. 18, 6520–6527 (2012).

38. Visser, E. J. et al. From Tethered to Freestanding Stabilizers of 14-3-3 Protein-Protein Interactions through Fragment Linking. Angew. Chem. Int. Ed Engl. 62, e202308004 (2023).

39. Wolter, M., et al. Fragment-Based Stabilizers of Protein–Protein Interactions through Imine-Based Tethering. Angew. Chem. Int. Ed. 59, 21520–21524 (2020).

40. Cossar, P. J. et al. Reversible Covalent Imine-Tethering for Selective Stabilization of 14-3-3 Hub Protein Interactions. J Am Chem Soc 143, 8454–8464 (2021).

41. Machleidt, T. et al. NanoBRET—A Novel BRET Platform for the Analysis of Protein–Protein Interactions. ACS Chem. Biol. 10, 1797–1804 (2015).

42. Dale, N. C., Johnstone, E. K. M., White, C. W. & Pfleger, K. D. G. NanoBRET: The Bright Future of Proximity-Based Assays. Front. Bioeng. Biotechnol. 7, (2019).

43. Robers, M. B. et al. Target engagement and drug residence time can be observed in living cells with BRET. Nat. Commun. 6, 10091 (2015).

44. Cho, E. J. & Dalby, K. N. Luminescence Energy Transfer–Based Screening and Target Engagement Approaches for Chemical Biology and Drug Discovery. SLAS Discov. 26, 984– 994 (2021).

45. Wade, M., Méndez, J., Coussens, N. P., Arkin, M. R. & Glicksman, M. A. Inhibition of Protein-Protein Interactions: Cell-Based Assays. in Assay Guidance Manual (eds. Markossian, S. et al.) (Eli Lilly & Company and the National Center for Advancing Translational Sciences, 2004).

46. Jiang, Z. et al. Adaptor-Specific Antibody Fragment Inhibitors for the Intracellular Modulation of p97 (VCP) Protein–Protein Interactions. J. Am. Chem. Soc. 144, 13218–13225 (2022).

47. Massoud, T. F. & Paulmurugan, R. Chapter 47 - Molecular Imaging of Protein–Protein Interactions and Protein Folding. in Molecular Imaging (Second Edition) (eds. Ross, B. D. & Gambhir, S. S.) 897–928 (Academic Press, 2021). doi:10.1016/B978-0-12-816386-3.00071-5.

48. Nadel, C. M., Ran, X. & Gestwicki, J. E. Luminescence complementation assay for measurement of binding to protein C-termini in live cells. Anal. Biochem. 611, 113947 (2020).

49. Xu, D., Farmer, A., Collett, G., Grishin, N. V. & Chook, Y. M. Sequence and structural analyses of nuclear export signals in the NESdb database. Mol. Biol. Cell 23, 3677–3693 (2012).

50. NanoBRET^TM^ Protein:Protein Interaction System Protocol. https://www.promega.com/resources/protocols/technical-manuals/101/nanobret-protein-protein-interaction-system-technical-manual/.

