## Supplementary Figures for "Development of a NanoBRET assay for evaluation of 14-3-3σ molecular glues"

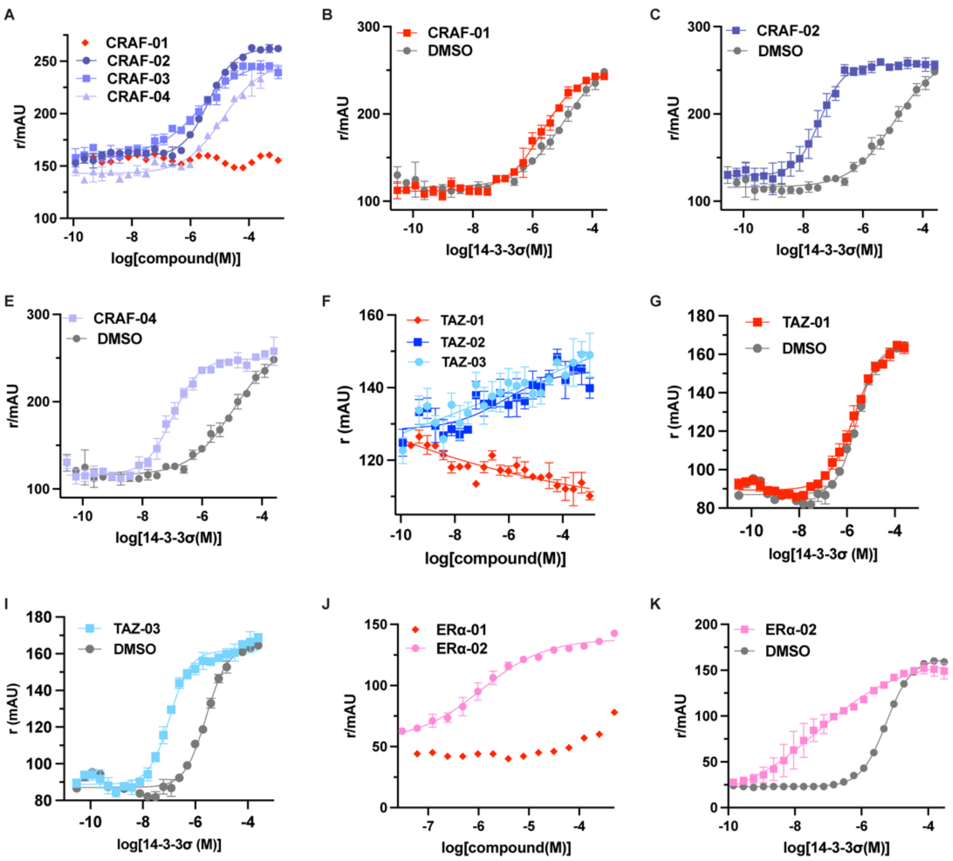


**Supplementary Figure 1:** CRAF pS259, TAZ pS89, and ERα pT594 peptide FA compound and protein titrations. **A)** CRAF-01-04 compound titrations. **B)** CRAF-01 protein titration. **C)** CRAF-02 protein titration. **D)** CRAF-03 protein titration. **E)** CRAF-04 protein titration. **F)** TAZ-01-03 compound titrations. **G)** TAZ-01 protein titration. **H)** TAZ-02 protein titration. **I)** TAZ-03 protein titration. **J)** ERα-01 and ERα-02 compound titrations. **K)** ERα-02 protein titration. All experiments n = 3.

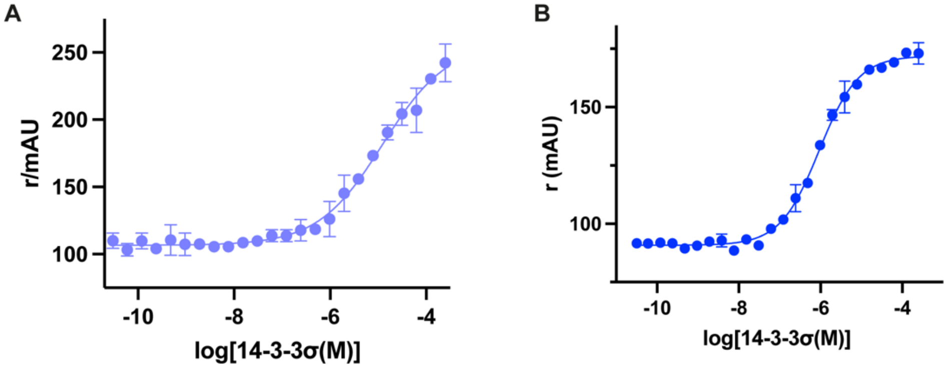


**Supplementary Figure 2:** 14-3-3/client peptide affinities. **A)** CRAF pS259 peptide FA $K_D$ assay (n = 3). **B)** TAZ pS89 peptide FA $K_D$ assay (n = 3).

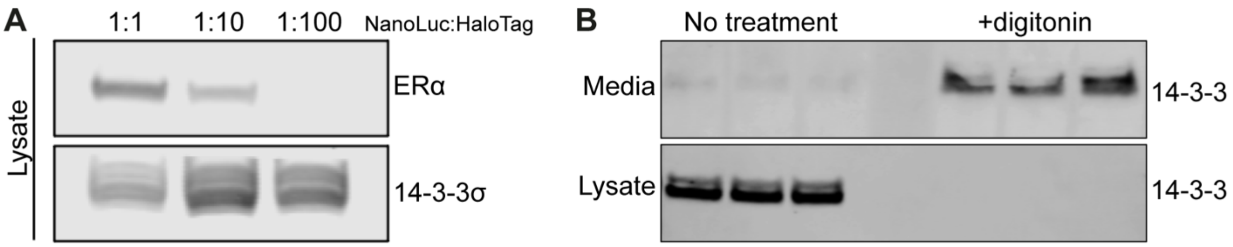


**Supplementary Figure 3:** Protein expression and effects of detergent. **A)** NanoLuc:HaloTag plasmid transfection ratios reflected in protein expression in HEK293T cells. **B)** Western blot for 14-3-3 without and with digitonin treatment (n=3). Cytosolic proteins are released into the media when treated when cells are treated with digitonin.

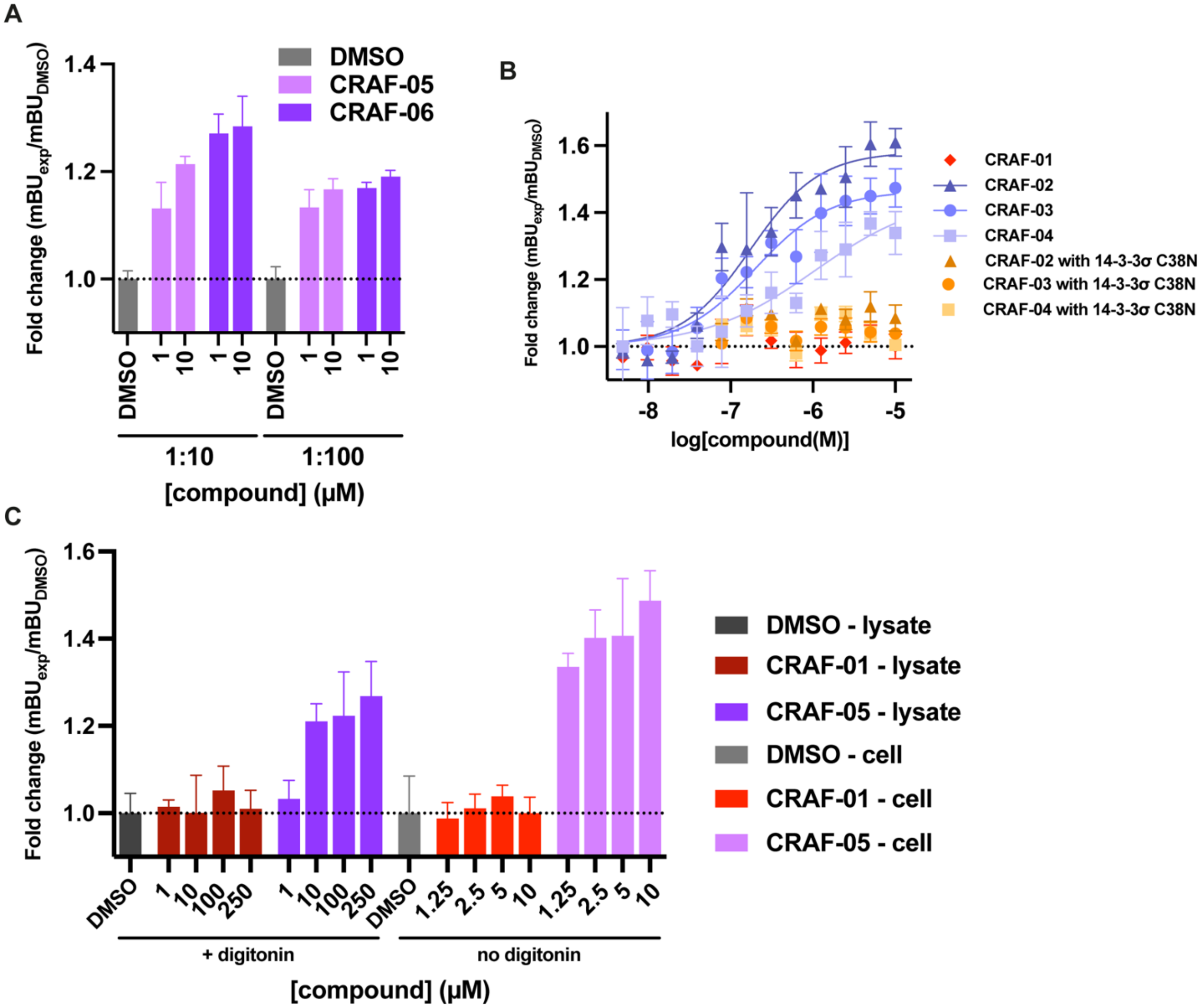


**Supplementary Figure 4:** CRAF/14-3-3σ NanoBRET optimization. **A)** CRAF/14-3-3σ 1:10 and 1:100 ratio comparison (n = 3). 1:10 resulted in a larger fold increase. **B)** 14-3-3σ WT vs 14-3-3σ C38N NanoBRET. Stabilizers were dependent on the presence of C38 (n = 3). **C)** Comparison of lysate and in cell NanoBRET assays (n = 3). Both showed stabilization with CRAF-05.

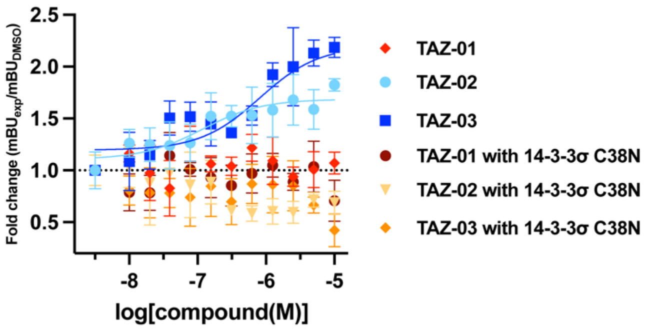


**Supplementary Figure 5:** TAZ NanoBRET with 14-3-3σ C38N (n = 3). Compounds bind to C38.

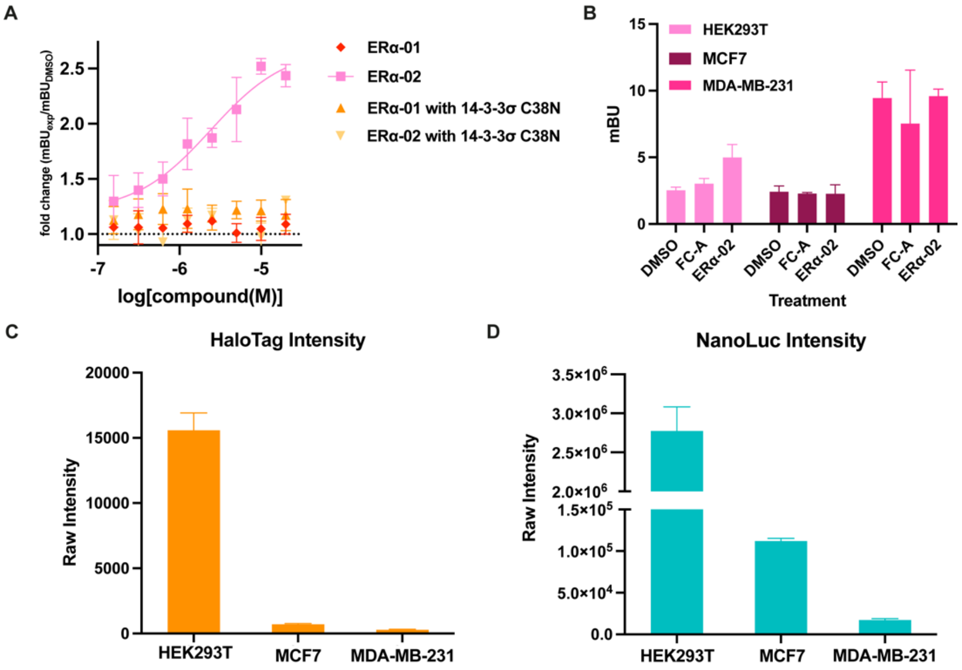


**Supplementary Figure 6:** ERα/14-3-3σ NanoBRET optimization. **A)** ERα NanoBRET with 14-3-3σ C38N (n = 3). Compounds are dependent on C38. **B)** ERα/14-3-3σ NanoBRET cell line comparison with FC-A (30 μM) and ERα-02 (10 μM) treatment (n = 3). HEK293T showed a measurable stabilization with ERα-02. **C)** Raw HaloTag luminescence intensity in HEK293T, MCF7, and MDA-MB-231 (n = 3). **D)** Raw NanoLuc luminescence intensity in HEK293T, MCF7, and MDA-MB-231 (n = 3).
